# Connexin 43 and Cell Culture Substrate Differentially Regulate OCY454 Osteocytic Differentiation and Signaling to Primary Bone Cells

**DOI:** 10.1101/2023.06.23.546276

**Authors:** Gabriel A. Hoppock, Evan G. Buettmann, Joseph A. Denisco, Galen M Goldscheitter, Sebastian N. Condyles, Otto J. Juhl, Michael A. Friedman, Yue Zhang, Henry J. Donahue

**Author notes:** Address correspondence to: Henry J Donahue, 70 S Madison St, Engineering Research Building, Rm 4332B, Richmond VA 23220. GAH and EGB contributed equally to this work.

## Abstract

Connexin 43 (Cx43), the predominate gap junction protein in bone, is essential for intercellular communication and skeletal homeostasis. Previous work suggests osteocyte-specific deletion of Cx43 leads to increased bone formation and resorption, however the cell-autonomous role of osteocytic Cx43 in promoting increased bone remodeling is unknown. Recent studies using 3D culture substrates in OCY454 cells suggest 3D cultures may offer increased bone remodeling factor expression and secretion, such as sclerostin and RANKL. In this study, we compared culturing OCY454 osteocytes on 3D Alvetex scaffolds to traditional 2D tissue culture, both with (WT) and without Cx43 (Cx43 KO). Conditioned media from OCY454 cell cultures was used to determine soluble signaling to differentiate primary bone marrow stromal cells into osteoblasts and osteoclasts. OCY454 cells cultured on 3D portrayed a mature osteocytic phenotype, relative to cells on 2D, shown by increased osteocytic gene expression and reduced cell proliferation. In contrast, OCY454 differentiation based on these same markers was not affected by Cx43 deficiency in 3D. Interestingly, increased sclerostin secretion was found in 3D cultured WT cells compared to Cx43 KO cells. Conditioned media from Cx43 KO cells promoted increased osteoblastogenesis and increased osteoclastogenesis, with maximal effects from 3D cultured Cx43 KO cells. These results suggest Cx43 deficiency promotes increased bone remodeling in a cell autonomous manner with minimal changes in osteocyte differentiation. Finally, 3D cultures appear better suited to study mechanisms from Cx43-deficient OCY454 osteocytes *in vitro* due to their ability to promote osteocyte differentiation, limit proliferation, and increase bone remodeling factor secretion.

**New and Noteworthy:** 3D cell culture of OCY454 cells promoted increased differentiation compared to traditional 2D culture. While Cx43 deficiency did not affect OCY454 differentiation, it resulted in increased signaling, promoting osteoblastogenesis and osteoclastogenesis. Our results suggest Cx43 deficiency promotes increased bone remodeling in a cell autonomous manner with minimal changes in osteocyte differentiation. Also, 3D cultures appear better suited to study mechanisms in Cx43-deficient OCY454 osteocytes.

## Introduction

Osteocytes are the most abundant cell type in mature bone, comprising approximately 90-95% of cells, and are important regulators of bone remodeling.(1, 2) These cells are terminally differentiated osteoblasts that develop after becoming embedded in newly secreted and mineralized bone matrix. Although once thought to be mainly dormant, increasing evidence shows osteocytes help maintain postnatal bone mass and quality via direct signaling to osteoblasts and osteoclasts. For example, osteocytes regulate osteoclast-mediated bone resorption by secreting the pro-resorptive cytokine receptor activator of nuclear factor-κB ligand (RANKL) to and its inhibitor osteoprotegerin (OPG).(3, 4) Furthermore, osteocytes regulate bone formation via the secretion of WNT inhibitors sclerostin and dickkopf-related protein 1 (DKK1) to osteoblasts.(5, 6) Aging, disuse, and estrogen deficiency—all associated with increased osteocyte apoptosis—show increased RANKL and sclerostin secretion.(7) Therefore, regulating osteocyte viability and paracrine signaling may be an effective way to preserve bone mass and alleviate osteoporosis resulting from aging, disuse, or estrogen deficiency.

One key protein heavily implicated in bone cell viability and signaling is connexin 43 (Cx43), the predominant gap junction protein in bone. Gap junctions are transmembrane protein channels that allow for cell-cell and cell-environment transfer of ions and small signaling molecules less than approximately 1 kDa. Gap junctions are comprised of two connexons, each comprising six transmembrane proteins called connexins. The importance of Cx43 in skeletal health and postnatal bone mass has been demonstrated in clinical and preclinical scenarios involving mutations in the *Gja1* gene, which encodes Cx43.(8) For example, humans with *Gja1* mutations develop oculodentodigital dysplasia and have craniofacial abnormalities, syndactyly, and decreased long bone mineral density and cortical thinning.(9–12)

Skeletal manifestations of *Gja1* mutations—such as cortical thinning and osteopenia—are also observed in murine models of conditional *Gja1* knockouts in osteoblasts and osteocytes. Deletion of *Gja1* in osteoblasts and osteocytes via late osteoblast stage non-inducible Cres (Col1 (2.3kb), osteocalcin, and DMP1 (8kb)) results in an osteopenic phenotype with decreased mechanical strength.(13–15) These results also suggest the observed osteopenic phenotype in mature bone arises due to enhanced osteocyte apoptosis and bone turnover with relative increases in endosteal bone-resorbing osteoclasts and periosteal bone-forming osteoblasts. Furthermore, these osteoclast and osteoblast changes are accompanied by alterations in signaling molecules such as RANKL, OPG and sclerostin within bone.(13) However, due to the widespread cellular targeting of DMP1 Cre, the cell-autonomous role osteocytic Cx43 plays in promoting this bone phenotype remains unknown.(16, 17)

Our knowledge of Cx43’s cell-autonomous role in osteocyte viability, differentiation, and signaling has been aided by immortalized osteocytic cell lines. There are several osteocytic cell lines used in research. Three of the most commonly used are MLO-Y4, IDG-SW3, and OCY454 cells, each of which have strengths and weaknesses.(18–21) This paper will focus on the most recently derived and available of these lines, the OCY454 cell line. This cell line was developed by the Pajevic Lab in 2015 from immortalized DMP1-GFP^+^ cells from murine long bones.(19) A major strength of this cell line is their ability to reliably express *Sost* (encodes sclerostin), without added differentiation factors, and at earlier timepoints than the MLO-Y4 and IDG-SW3 lines (< 3 weeks). For example, Spatz et al. demonstrated *Sost* expression was detectable in as little as 3 days of differentiation when OCY454 cells were cultured on Alvetex Reprocell 3D scaffolds, which better recapitulate the bone microenvironment.(19) Therefore, OCY454 cells in 3D microenvironments may be ideally suited to study the role of Cx43 deficiency on mature osteocyte biology.

The genetic deletion or pharmacological inhibition of Cx43 has been studied in both MLO-Y4 and IDG-SW3 osteocyte cell lines, but only in 2D culture conditions. These studies demonstrated Cx43 deficiency in either cell line led to increased osteocyte resorptive signaling (RANKL/OPG) and osteoclast differentiation, similar to *in vivo* reports.(8, 13, 14, 22) Furthermore, Cx43 deficiency in IDG-SW3 cells demonstrated impaired differentiation into mature osteocytes as evidenced by attenuated Dentin matrix protein-1 (*Dmp1*) and *Sost* expression.(8) However, no studies to date have investigated how the cellular microenvironment and Cx43 deficiency both modulate OCY454 osteocyte proliferation, differentiation, and cell signaling to bone effector cells (osteoblasts and osteoclasts). To evaluate this, OCY454 cells were made Cx43 deficient by CRISPR-Cas9 genetic engineering and cultured on collagen type 1 coated 2D tissue culture plastic substrates (TCPS) or 3D Reprocell (Alvetex™) scaffolds. First, OCY454 wildtype (WT) and Cx43 deficient (Cx43 KO) cells’ proliferation and differentiation were probed by molecular analysis when seeded in 2D or 3D (**Figure 1A**). Then, conditioned media from OCY454 cells was used to determine the role of Cx43 and culture microenvironment in regulating osteocyte signaling to osteoblasts and osteoclasts (**Figure 1B**). We hypothesized in both 2D and 3D cultures, Cx43 KO cells would show delayed differentiation and pro-osteoclastic and pro-osteoblastic signaling compared to their WT counterparts. We also hypothesized 3D cultures would promote earlier differentiation compared to 2D cultures due to their ability to mimic the native osteocyte environment more accurately.

**Figure 1.**
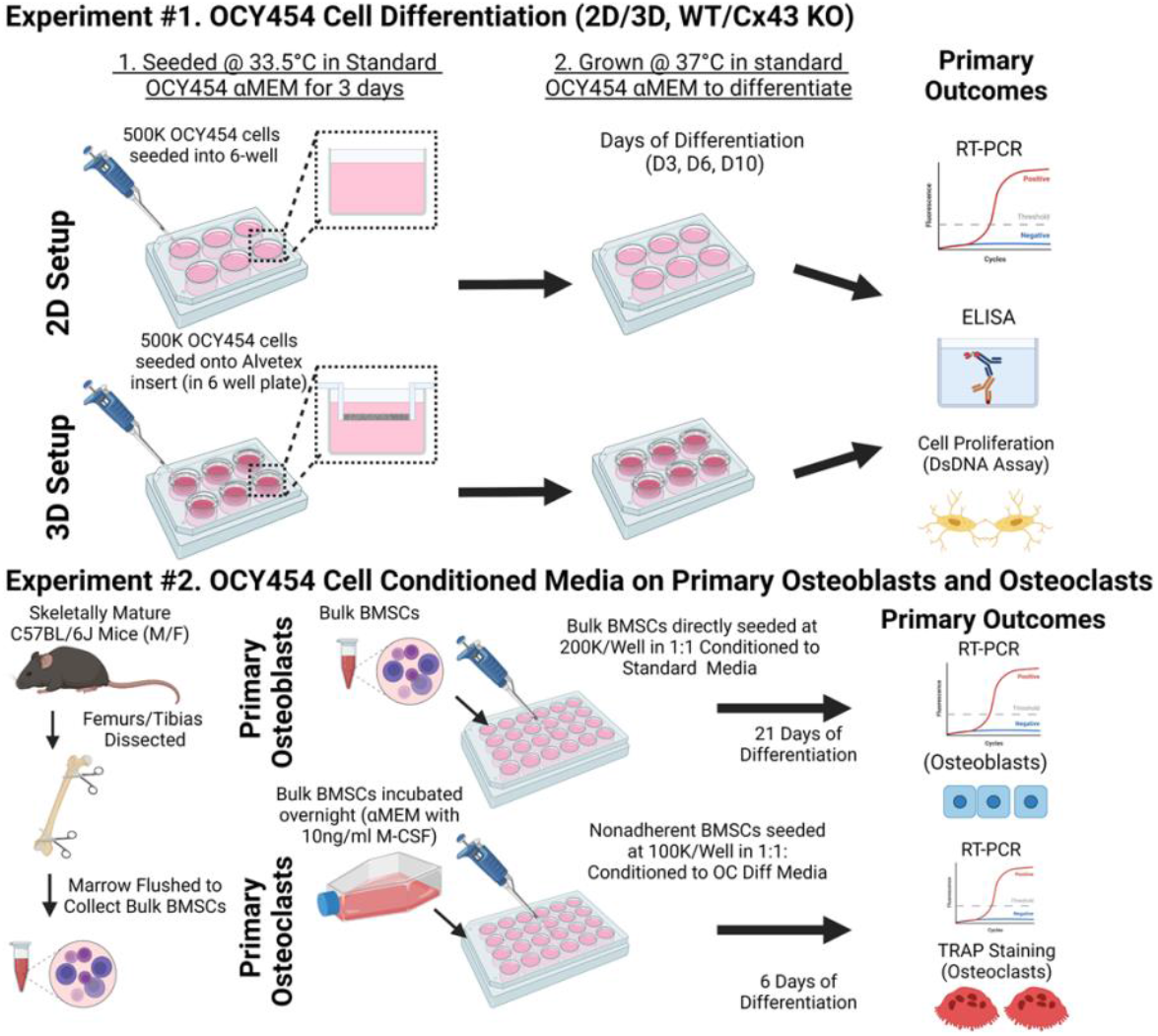
Timeline and Experimental Protocol for Study. A. OCY454 osteocyte-like wildtype (WT) and Cx43-deficient (KO) cells were cultured in Collagen1 coated plastic (TCPS - 6 well) or Alvetex (Reprocell 12 well inserts) at 500,000 cells/well for 2D and 3D culture, respectively. Cells were cultured in 10mL standard OCY454 αMEM media for 3 days at 33.5°C before switching to 37°C to induce differentiation. OCY454 WT and KO cells (on 2D/3D) were harvested for DNA, RNA, and protein at 3, 6, or 10 days post-differentiation (D3, D6, D10). **B:** Adult bone marrow stromal cells (BMSCs) were collected from adult C57BL/6J mice (16-20 weeks old: male and female cells pooled) and directly seeded at 200,000 cells/well (TCPS 24-well) for primary osteoblast assays. Osteoblast differentiation was measured by osteoblastic gene expression following 21 days of culture in 50% fresh OCY454 standard media:50% conditioned media. For osteoclastogenesis, non-adherent BMSCs were seeded at 100,000 cells/well (TCPS 24-well) in 50% conditioned media 50% : 50% Osteoclast Differentiation Media (Standard OCY454 αMEM supplemented with 25 ng/ml M-CSF, 5 ng/ml RANKL). Positive controls consisted of growth in standard OCY454 αMEM supplemented with 25 ng/ml M-CSF and 25 ng/ml RANKL. Following 6 days of culture, osteoclast differentiation was measured by TRAP stained cell quantification and osteoclastic gene expression. Figure created with BioRender.

## Methods

### Connexin 43 Deletion

OCY454 cells were kindly provided by the laboratory of Paola Pajevic and have been previously characterized.(19) An OCY454 Cx43-deficient cell line was generated by CRISPR*-*Cas9 system at VCU Biological Macromolecule Shared Resource. Briefly, a single guide RNA sequence (AACCCTACCCCCCCA) was designed to generate a frameshift mutation approximately 20 amino acids downstream of the start ATG of Cx43 and cloned into the guide RNA vector described by the Church laboratory.(23) The guide RNA vector was co-transfected into OCY454 cells together with humanized Cas9 and a puromycin resistance cassette. Following 48 hr of selection for puromycin resistance, single cells were dilution cloned into collagen coated 96-well plates and expanded for 10 days at 33°C. Viable clones were screened for the presence of mutations using the Surveyor Mismatch Endonuclease Assay (Integrated DNA Technologies), followed by DNA sequencing to validate the presence of nonsense mutations generated by frame shift. Western Blot and parachute assay were further utilized to verify a decrease in Cx43 protein levels (**Figure 2A**) as well as decreased gap junctional intercellular communication (**Figure 2B**). Both wildtype and connexin 43 deficient OCY454 cells were expanded multiple passages (passages 10-12) and then stored in -80°C in BAMBANKER™ (LYMPHOTEC Inc.) until further use.

**Figure 2.**
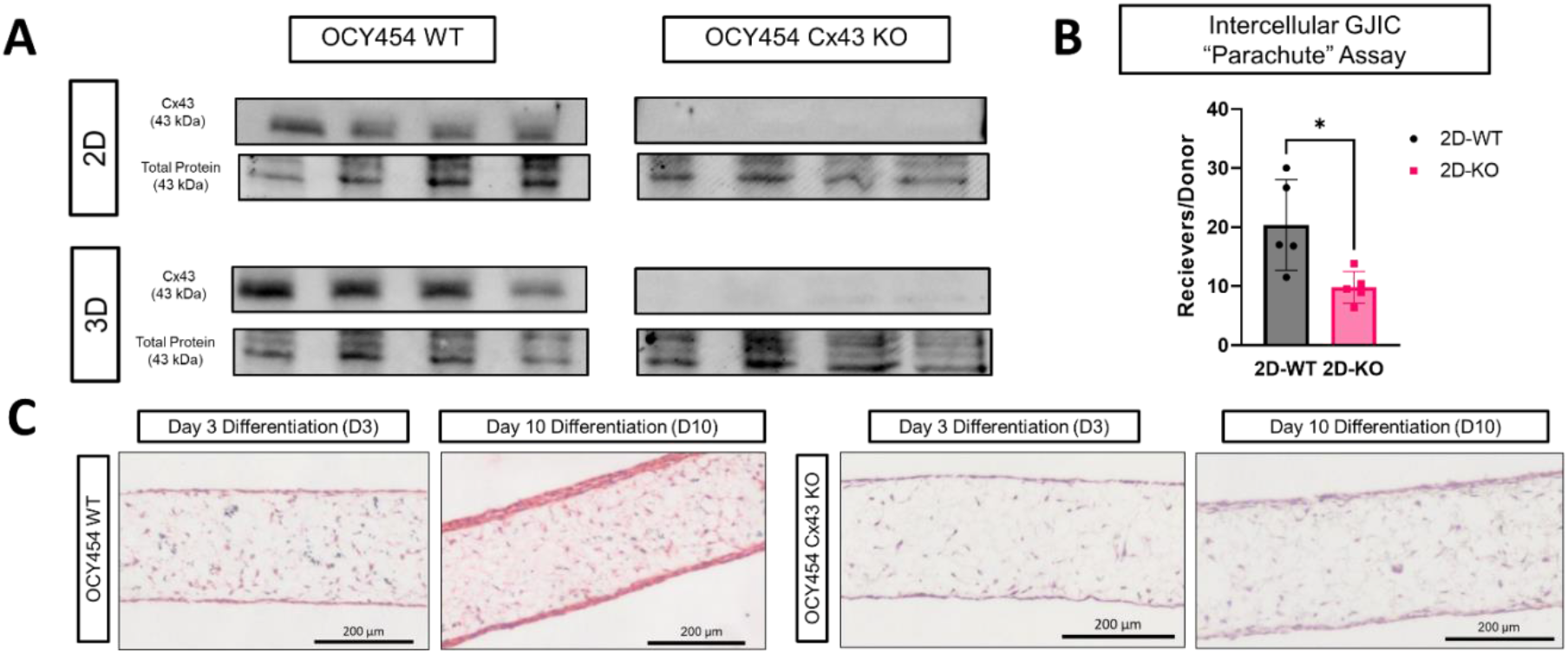
Cx43 protein was deleted from OCY454 Cells using CRISPR_Cas9 and resulted in impaired GJIC. Panel A. OCY454 osteocyte-like wildtype (WT) and Cx43-deficient (KO) cells differentiated for 10 days on 2D and 3D substrates. OCY454 WT showed Cx43 protein staining on western blot, but not Cx43-deficient (KO) cells. **Panel B:** OCY454 Cx43 KO cells had significantly less Calcein Blue AM transfer. **Panel C:** Cross-sections of Hematoxylin and Eosin Y stained 3D Alvetex scaffolds (200 micron thickness) following 3 and 10 days of differentiation of OCY454 osteocyte-like wildtype (WT) and Cx43-deficient (KO) cells. OCY454 cells in both genotypes are seen to infiltrate the interior and edges of the entire scaffold. Despite equal seeding densities, Cx43 KO cells appear to infiltrate and cover these scaffolds to a lesser degree. Scale bar is 200 microns. * p < 0.05

### OCY454 Cell Culture

Both WT and Cx43 KO OCY454 cells following thawing were seeded on rat type I collagen-coated plates and grown at 33° C in standard OCY454 media (**Supplemental Table 1**: αMEM with 10% FBS (Gibco: 26140-079; Lot # 2496659RP) and 2% P/S). After reaching 80-90% confluence, OCY454 cells were trypsinized and seeded onto either rat-tail collagen-coated (100µg/ml; Corning: CLS354236; Lot # 2195003) 6-well plates or 3D polystyrene scaffolds (Alvetex™ 12-well inserts: Reprocell AVP005) at a density of 500,000 cells/well or scaffold. Cells were cultured with standard media at 33°C for 3 days for proliferation (permissive temperature for SV40 Large T Antigen) and were then moved to 37°C (semi-permissive temperature) for 3-10 days to promote differentiation (D3-D10) before harvesting for subsequent DNA, RNA, or protein analysis of media and cell lysate. Standard media was changed 3X weekly (Monday, Wednesday, Friday) and 24 hours prior to conditioned media harvest. In total, there were 4 distinct experimental groups used: WT cells on TCPS (2D-WT), KO cells on TCPS (2D-KO), WT cells on Alvetex scaffolds (3D-WT), and KO cells Alvetex scaffolds (3D-KO).

### Validation of Connexin 43 Deletion (Western Blot and Dye Transfer Assay)

Total protein from OCY454 cell (WT and KO) lysate was collected in duplicate from TCPS 6-well plates or Alvetex 12-well inserts following ten days of differentiation at 37°C by RIPA Buffer (Thermo Fisher: 50-152-132) containing 1% proteinase inhibitor (Sigma: P8340). Protein concentrations were assayed by BCA assay (Thermo Fisher: 23227) according to the manufacturer’s protocols. For western blots, 50 ug of protein was prepared in 4X Laemmli-Buffer (Bio-Rad) before being separated on 4-20% Tris Glycine PAGE gel (200V; 40 mins). Next protein was transferred to PVDF membrane using Trans-Blot turbo (Bio-rad) and blocked for 60 mins in Odyssey Blocking Buffer with TBS (LI-COR Biosciences #927-50010). Blots were incubated with rabbit anti-mouse Cx43 antibody (Sigma C6219; Lot # 018M4842V; 1:8000 dilution) overnight with gentle agitation at 4°C. The next day blots were washed, incubated with anti-rabbit HRP antibody (Cell Signaling Technology 7074P2; 1:3000 dilution) for 1 hour at room temperature and developed with ECL substrate solution (Bio-Rad #1705060).

To verify Cx43 deletion led to altered gap junction intercellular communication (GJIC), cells underwent the dye coupling parachute assay as previously described.(24) WT and KO cells were cultured in 6-well plates as previously described (in *OCY454 Cell Culture*) until D0 of differentiation. At D0 of differentiation, hFOBs-1.19 (hFOB; human fetal osteoblasts) donor cells were simultaneously labeled with 10 μM Calcein Blue-AM (transferrable through GJIC) and 10 μM DiI solution (nontransferable through GJIC) for 30 mins. hFOBs were used as donor cells due to their large dye carrying capacity and lack of endogenous fluorescent transgenes. Labeled hFOBs were then trypsinized, centrifuged at 200 g for 5 min, and counted. Labeled hFOB cells were then dropped (parachuted) onto 90-100% confluent plates of unlabeled 2D-WT and 2D-KO cells at a 1:500 ratio of labeled hFOB (∼2500 cells) to unlabeled OCY454 cells (∼1 million cells) for 2 hours at 37°C. Mixed cell populations (donor + receiver) were then trypsinized and fixed with 2% paraformaldehyde for 15 min. Samples were washed and resuspended in FACS tubes for cell sorting. Pure donor hFOBs and un-treated OCY454 receiver cells served as positive and negative controls, respectively. The first 10,000 cell events were counted, with ratio of receiver (Calcein Blue AM^+;^ DiI^−^) to donor (DiI^+^) computed and analyzed as performed previously.(25)

### Cell Count

Prior to harvest, cells were washed twice in sterile PBS to remove dead cells and debris. DNA was harvested from cells according to manufacturer standards using the Quant-iT™ PicoGreen™ dsDNA Assay Kits (Thermofisher, Catalog # P11496). OCY454 cells were collected directly from scaffold or well plate using 1 mL of TE lysis buffer (10 mM Tris, ph8, 1 mM EDTA and 0.2% (v/v) Triton X-100). Samples were incubated for 30 mins at 4°C with occasional vortexing to ensure lysis before centrifugation (10,000g for 15 mins) and collection of supernatants. For the assay, 10uL of supernatant were diluted to 100ul in 1X TE buffer and added to flat-bottom 96-well plate. 100 uL of PicoGreen™ reagent was added to each well, incubated for 5 mins, and the resultant fluorescence intensity (FI) per well was read at 460 nm and 540 nm (excitation and emission) on a plate reader (Perkin-Elmer; 500 ms exposure). FI results were correlated to cell numbers based on a 2-fold dilution standard curve prepared from WT and KO cells counted with a hemacytometer.

### Histology

Standard paraffin sectioning with hematoxylin and eosin staining was used to visualize OCY454 WT and KO cellular distribution within 3D Alvetex scaffolds following day 3 and day 10 of differentiation (D3 – D10). At harvest, multiple 3D Alvetex scaffolds were washed in PBS and then partitioned into 3mm diameter inserts via biopsy punch (Millitex Catalog # 33-32-P/25). These 3mm diameter inserts were fixed in 4% PFA overnight and subsequently processed according to manufacturing protocols (Reprocell Inc., Beltsville, MD) for paraffin sectioning. The 3mm inserts went through a dehydration series, followed by infiltration with HistoChoice® Clearing Agent (Sigma-Aldrich Catalog #H2779), and paraffin embedding. Sagittal sections (5-7 microns) were cut on a microtome, rehydrated and stained for hematoxylin and eosin according to manufacturer protocols (Reprocell Inc., Beltsville, MD). Slides were cover-slipped with DPX mounting medium (EMS Catalog #13510) and imaged at 20X using a Zeiss Inverted Microscope and stitched together in ZenPro.

### Gene Expression

Prior to harvest, cells were washed twice in sterile PBS to remove dead cells and debris. Cells were lysed and RNA was extracted and purified based on manufacturer protocols (Qiagen RNeasy Kit #74106). 600uL of RNA lysis Buffer (with 1% (v/v) β-mercaptoethanol) was used to lyse cells, RNA was collected and purified, with on-the-column genomic DNA removal performed with the RNase-Free DNase Set (Qiagen #79256). RNA concentration was quantified by NanoDrop Lite (Thermo Fisher #ND-LITE), and RNA was stored at -80°C. RNA quality was spot-checked by Bioanalyzer (Agilent # 5067-1512) and always exceeded RNA Integrity Numbers (RINs) of 9 out of 10, demonstrating minimal degradation.(26) For cDNA generation, 500ng of RNA was reverse transcribed into cDNA using the Biorad iSCRIPT kit (Biorad #1708841). SYBR (Thermo Fisher #A25742) based real time quantitative PCR (RT-qPCR) with 10 ng RNA equivalent cDNA per reaction in triplicate to assess relative abundance and amplicon specificity. RT-qPCR was also performed on RNA samples without reverse transcription (NRT) to assess genomic DNA contamination. Primer sequences for OCY454 cells were from the BioRad PrimePCR catalog (**Supplemental Table 2**) and included genes implicated in early osteocyte differentiation (*Mmp14*; encodes matrix metallopeptidase 14, *Pdpn*; encodes podoplanin, *Dmp1*; encodes dentin matrix acidic phosphoprotein 1), proliferation (*Ccnd1*; encodes cyclin D1), apoptosis (*Ddit3*; encodes DNA damage inducible transcript 3), WNT/β-catenin signaling (*Sost*; encodes sclerostin, *Axin2*; encodes axis inhibition protein 2, *Ctnnb1*; encodes β-catenin) and bone metabolism (*Tnfsf11*; encodes receptor activator of nuclear factor kappa-? ligand (RANKL), *Tnfrsf11b*; encodes osteoprotegerin).(1) A custom primer for an additional marker of OCY454 differentiation, *Phex* (encodes phosphate regulating endopeptidase x-linked), was used with sequence previously reported (Forward 5’-3’: GAAAGGGGACCAACCGAGG; Reverse 5’-3’: AACTTAGGAGACCTTGACTCAC).(19) Relative gene expression was assessed for each sample by normalization to the reference gene *Actb* (encodes β-actin).

### Conditioned Media and ELISAs

Twenty-four hours prior to conditioned media harvest, cells were replenished with fresh standard OCY454 media (**Supplemental Table 1**). Conditioned media for ELISA and primary cell experiments were centrifuged at 500g for 5 mins at 4°C to remove cells and debris and frozen at -80°C until use. Two mL of conditioned media was collected from each well and aliquoted for ELISA. The remaining conditioned media for all samples (n > 6 per group) was pooled per experimental group (2D/3D and WT/KO) and used for primary cell differentiation experiments. Aliquots of individual OCY454 WT and Cx43 KO samples differentiated for ten days were assessed for key secretory proteins implicated in osteoblast and osteoclast differentiation – sclerostin (R&D Systems; Catalog #DY1589-05) and dickkopf-related protein 1 (R&D Systems; Catalog #DY1756). All ELISA assays were run in duplicate according to manufacturer protocols, supplied standards and controls ELISA (R&D Systems; DuoSet Ancillary Kit #2 Catalog #DY008). Standards were fit to 4 parameter logistic curves using GraphPad Prism Pro (Version 8; La Jolla, CA) following optical density readings at 450 nm, with correction wavelength 540 nm.

### Primary Cell Culture

All animal procedures were approved by VCU IACUC (Protocol # AD10001341). Skeletally mature C57BL/6J mice (16-20 weeks) were used for all primary cell culture with pooling of bone marrow stromal cells (BMSCs) from the long bones (tibias/femurs) of an equal number of male and female mice (n = 2-3 mice per sex per harvest). Primary BMSCs were prepared for osteoblast and osteoclast conditioned media experiments according to previous methods.(27) In brief, mice were euthanized by CO2 narcosis followed by cervical dislocation and long bones were aseptically harvested. Long bone marrow was collected by centrifugation (10,000g), resuspended in standard OCY454 media (**Supplemental Table 1**) and passed through a cell strainer (40µm nylon mesh; BD Biosciences) and counted before use in primary osteoclast and osteoblast experiments.

### Osteoblast Conditioning Experiment

Following primary bone marrow cell isolation, cells were directly plated at 200,000 cells/well of 24-well-plates and grown at 37°C (5% CO2) with 50% OCY454 conditioned: 50% standard OCY454 media to promote osteoblastogenesis. Conditioned media came from pooled samples of OCY454 cells differentiated over 10 days from one of four experimental groups: 2D-WT, 2D-KO, 3D-WT, 3D–KO to eliminate inter-specimen variability. Osteoblast positive control media (OB PC media; **Supplemental Table 1**) consisted of standard media supplemented with 10mM β-glycerophosphate (Sigma Catalog # G9422; Lot # SLBN8374V) and 50 ug/ml ascorbic acid (Sigma Catalog # A4403; Lot # LEK1177). Osteoblast negative control media samples contained only standard OCY454 media. Primary BMSCs were cultured under these conditions at 37°C (5% CO_2_) for 3 weeks to promote osteoblastogenesis with media changes twice per week. At day 21, cells were lysed for RNA (Qiagen RLT lysis buffer). Relative gene expression of key osteoblast differentiation (*Runx2;* encodes Runt-Related Transcription Factor 2) and bone formation markers (*Alpl;* encodes Alkaline Phosphatase and *Col1a1*; encodes Collagen Type I Alpha 1 Chain) was assessed for each sample by normalization to the reference gene *Actb*.

### Osteoclast Conditioning Experiment

Following primary bone marrow cell (BMSC) isolation, BMSCs were plated in T-75 flasks overnight with standard OCY454 media supplemented with 10ng/ml M-CSF (R&D Systems Catalog #416-ML-010) at 37°C in 5% CO2. Non-adherent bone marrow cells were collected and seeded at 100,000 cells/well of 24 well-plates and grown at 37°C (5% CO2) with 50% OCY454 conditioned media: 50% osteoclast differentiation media (OC Diff Media; **Supplemental Table 1**) to promote osteoclastogenesis. Osteoclast differentiation media consisted of standard OCY454 media supplemented with 25ng/ml M-CSF (R&D Systems Catelog # 416-TEC; Lot # ME5021121) and 5 ng/ml RANKL (R&D Systems Catalog #462-TEC-010; Lot # CWA2622011), which is the previously reported EC_50_ needed for primary bone marrow cell differentiation into osteoclasts. Conditioned media came from pooled OCY454 cells differentiated for 10 days from one of four experimental groups: 2D-WT, 2D-KO, 3D-WT, 3D-KO. Osteoclast positive control media (OC PC media; **Supplemental Table 1**) samples consisted of standard αMEM supplemented with 25ng/ml M-CSF and 25ng/mL RANKL. Osteoclast negative control media (OC NC media; **Supplemental Table 1**) samples contained only 10ng/mL M-CSF and no RANKL. Primary nonadherent BMSCs were cultured under these conditions at 37°C (5% CO2) for 6 days, when large multinucleated cells were present in OC PC media condition, with media changes twice before harvest. At harvest, cells were fixed overnight with 4% PFA for TRAP staining or immediately lysed for RNA isolation. TRAP positive cells (TRAP^+^) were stained and quantified according to previous reports.(27) Specifically, five random 10x magnification fields per well were imaged under brightfield settings and averaged for quantitative outcomes. TRAP^+^ images were analyzed using particle analysis in FIJI software suite (Threshold: 120; >500 pixels^2^, circularity 0-1) to assess average TRAP^+^ cell size per field. Relative gene expression of key osteoclast resorption marker (*Ctsk;* encodes cathepsin K) was assessed for each sample by normalization to the reference gene *Actb*.(28)

### Statistics

Statistical analyses were performed using GraphPad Prism (Version 9.3) software. All data are expressed as means ± standard deviation with individual data points for each column representing total sample size. Data were analyzed using two-way analysis of variance (ANOVA) or three-way ANOVA with Tukey’s post-hoc or by student’s t-test, as appropriate. For all comparisons, p < 0.05 was considered significant.

## Results

*CRISPR-Cas9 Deletion of Gja1 Gene in OCY454 Cells Led to No Detectable Cx43 Protein Production and Decreased Gap Junction Intercellular Communication (GJIC)* Western blot analysis for Cx43 protein on collagen coated 2D-WT and 3D-WT cells cultured for ten days revealed strong Cx43 staining (**Figure 2A; at 43 kDa**). In contrast, puromycin resistant 2D-KO and 3D-KO cells cultured simultaneously demonstrated no discernable Cx43 band at 43 kDa, thereby validating CRISPR-Cas9 mediated protein deletion (**Figure 2A**). To determine if this loss of Cx43 in OCY454 cells affected GJIC, a dye transfer assay previously used by our group was used.(25) Donor hFOBs containing fluorescent permeable (Calcein Am – Blue) and impermeable (DiI) dyes were parachuted onto confluent 2D-WT and 2D-KO cells to assess gap junction communication. Following 2 hours of incubation of hFOBs with OCY454 cells, there was significantly greater permeable Calcein Blue Am dye transfer in 2D-WT versus 2D-KO cells. On average, 20.38 ± 7.67 neighboring 2D-WT cells received dye transfer from donor cells compared to 9.82 ± 2.70 neighboring KO cells (**Figure 2B;** p < 0.05). These results suggest a 50% decrease in GJIC in 2D-KO cells compared to 2D-WT. H&E histology on 3D scaffolds demonstrated that OCY454 cells in both genotypes infiltrated the interior and edges of the entire scaffold (**Figure 2C**). In all, these data suggest successful incorporation of OCY454 onto 3D Alvetex scaffolds and deletion of Cx43 in OCY454 cells, leading to decreased, but not absent, GJIC.

*Cell Culture Substrate and Genotype Differentially Regulate OCY454 Cellular Proliferation* To examine changes in cellular number due to cell culture substrate and Cx43 deficiency, we quantified dsDNA following cell lysis during D3-D10 of OCY454 cell differentiation. By D10, the cell count (derived from DNA standard curve) of WT and KO cells on both substrates (2D and 3D) was significantly increased compared to D3 of differentiation (**Figure 3A**). This trend was also observed on the 3D samples via H&E staining comparing D3 to D10 (**Figure 2C**). While cell count increased in all groups at D10 compared to D3, genotype only affected cellular proliferation in 3D cultures. For example, with 2D culture substrate, both 2D-WT and 2D-KO cell counts continued to increase throughout the 10-day course of the experiment, regardless of genotype (**Figure 3A**, genotype p = 0.2548). In contrast, OCY454 cells cultured on 3D showed a significant main interaction due to genotype and timepoint (Figure 3A; p < 0.0001). Specifically, while we observed 3D-WT and 3D-KO cells had similar cell numbers at D3, there were significantly more 3D–WT cells compared to 3D–KO cells at later timepoints (D6 and D10; **Figure 3A**).

**Figure 3.**
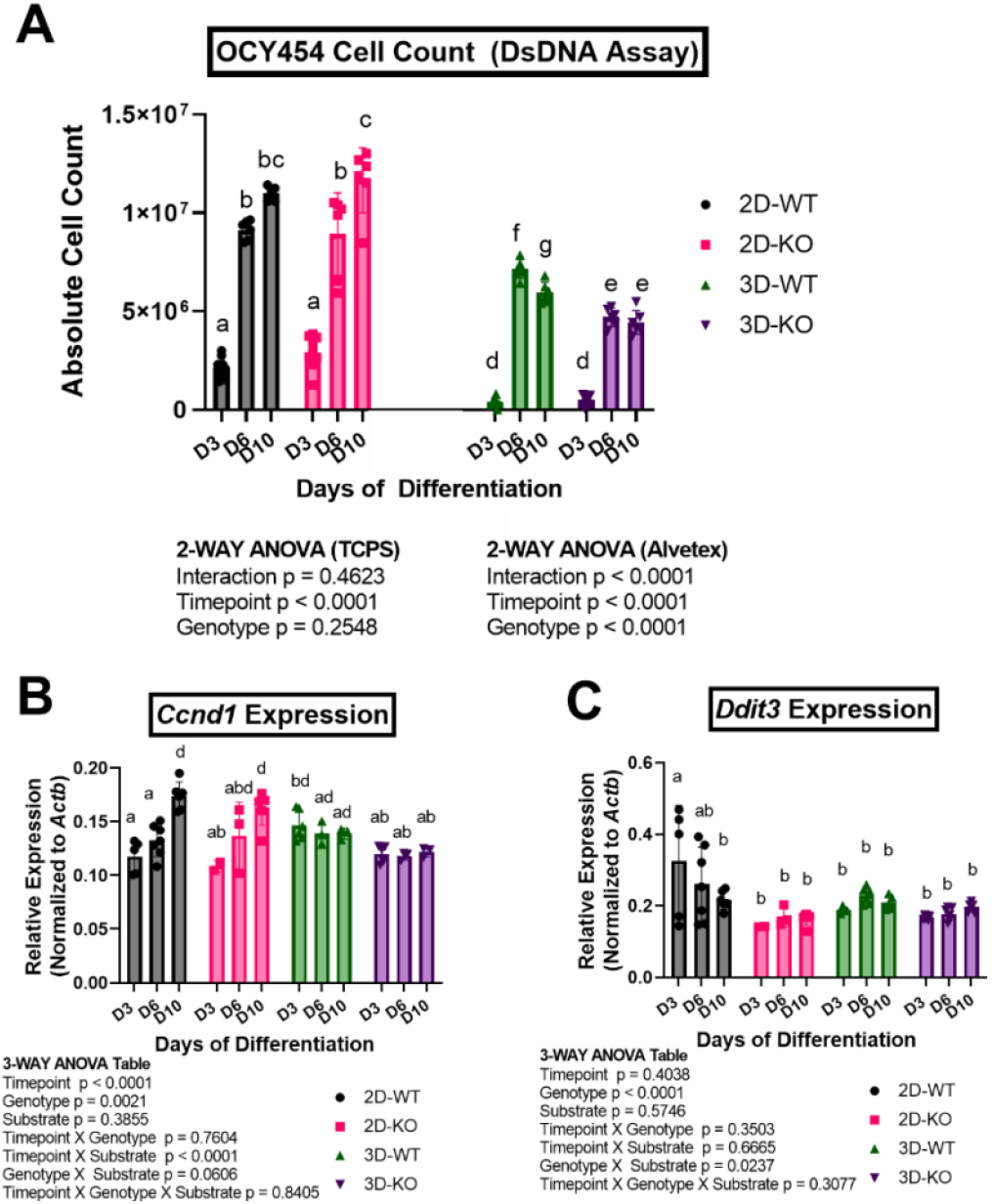
Culture of OCY454 cells on 2D lead to greater cell number over time compared to 3D. Cx43 deficiency lead to decreased OCY454 cell number compared to WT cells on 3D only. Panel A. Changes in OCY454 osteocyte-like wildtype (WT) and Cx43-deficient (KO) cells grown on either tissue culture plastic (2D) or Alvetex (3D) were computed throughout differentiation by Picogreen DsDNA fluorescent assay. Absolute cell counts for 2D and 3D samples weren’t directly compared via statistics due to different overall growth areas. **Panel B:** Relative *Ccnd1* expression (encodes cyclin D1), suggested significantly increasing rates of proliferation over time (D10 versus D3) on 2D but not 3D cell culture formats (both genotypes). **Panel C:** Relative *Ddit3* expression (encodes DNA damage inducible transcript 3), suggested significantly increased rates of apoptosis overall in 2D-WT cells at D3 compared to other genotypes and timepoints Columns not sharing a letter are significantly different via Tukey’s post-hoc test following applicable 3-Way or 2-Way ANOVA (main effects: substrate, genotype, timepoint), p<0.05. Data is presented mean ± SD with data points for each column representing total n

We also examined *Ddit3* and *Ccnd1* expression to determine if changes in cell number were due to cellular apoptosis or cellular proliferation, respectively. *Ccnd1* expression was increased due to timepoint (**Figure 3B** p < 0.0001) and genotype (**Figure 3B**; p = 0.0021), with D10 and WT cells, respectively, showing the most *Ccnd1* expression. Furthermore, *Ccnd1* demonstrated a significant main interaction with timepoint x substrate (**Figure 3B**; p < 0.0001). For example, 2D cells demonstrated increased *Ccnd1* expression over time (**Figure 3B**) while 3D cultured cells showed no changes in *Ccnd1* expression over time (**Figure 3B**).

Gene expression data revealed significantly increased *Ddit3* expression in the 2D-WT cells, but only at D3, suggesting increased cellular apoptosis in this group compared to other genotypes and substrates (**Figure 3C**; Genotype x Substrate Interaction p = 0.0237). All other groups (2D–KO, 3D–WT, and 3D– KO) demonstrated similar levels of *Ddit3* expression to each other and over time, suggesting constant rates of apoptosis. In all, these data suggest an increased number of OCY454 cells when cultured on 2D compared to 3D (regardless of genotype) likely due to greater proliferation on 2D vs 3D. In contrast, OCY454 cells cultured on 3D had minimal changes in cell number from D6 onward, which are reflective of the steady levels of *Ccnd1* and *Ddit3* expression at this time.

*3D Substrate Promotes Increased Expression of Osteocytic Maturation Markers Compared to 2D Substrate; Connexin 43 Deficiency Causes Minimal Decrease in Osteocyte Markers of Differentiation* To examine the maturation of the OCY454 cells due to substrate (2D vs. 3D) and Cx43 deficiency (WT vs. KO), we examined gene expression changes over the time course of differentiation (D3-D10) using well-established markers of osteocyte differentiation.(1) Examining expression of early markers of osteoblast-osteocyte transition (*Mmp14, Pdpn*), revealed significant main effects of culture substrate and timepoint but not genotype (**Figure 4A**; p < 0.05). Expression of *Mmp14* was similar on 2D over time but significantly increased on 3D over the course of differentiation regardless of genotype (**Figure 4A**). Culture substrate modulated expression of *Pdpn* over time, as its expression decreased on 2D and increased on 3D when comparing D3 to D10, regardless of genotype (**Figure 4A**). The later osteocyte maturation markers *Phex* and *DMP1* showed significantly increased effects of timepoint and substrate (**Figure 4B**; p < 0.0001), with more subtle but significant interactions of genotype and timepoint (**Figure 4B**; p < 0.05). For example, *Dmp1* and *Phex* expression on 3D-WT and 3D-KO significantly increased over time compared to 2D–WT and 2D-KO counterparts, which decreased or remained stagnant in 2D (**Figure 4B**). Specifically, 2D-KO cells showed a significant decrease in *Dmp1* expression at D10 compared to their 2D-WT counterparts, however, this was not seen at other timepoints with our 2D or 3D samples. *Sost and Dkk1* expression, late markers of osteocyte maturation, were only detected on 3D samples (**Figure 5**). However, *Sost and Dkk1* expression were not significantly different between 3D-WT and 3D-KO samples (**Figure 5A**; p > 0.05). Overall, increased expression of osteocyte maturation markers over time in 3D versus 2D samples suggest enhanced osteocyte differentiation using a 3D culture substrate. In addition, only *DMP1* expression was significantly affected by genotype and demonstrated decreased expression in 2D-KO versus 2D-WT cells only at D10. Therefore, Cx43 deficiency did not significantly inhibit OCY454 cell differentiation based on the gene expression of multiple well-defined osteocyte markers.

**Figure 4.**
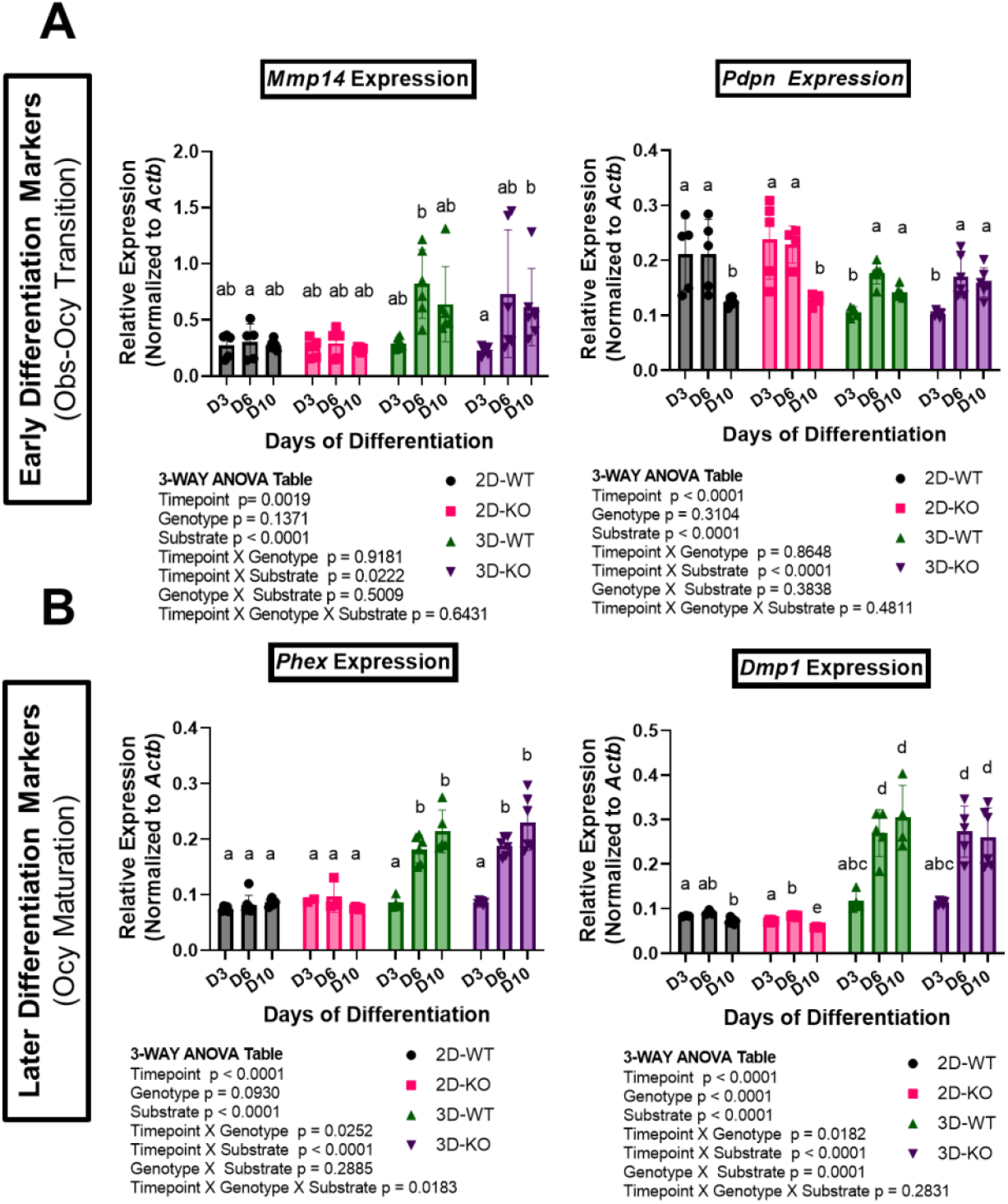
2D culture led to impaired markers of osteocyte maturation in OCY454 cells compared to 3D culture. Cx43 deficiency had a minimal effect on expression of osteocyte differentiation markers. Changes in differentiation of OCY454 osteocyte-like wildtype (WT) and Cx43-deficient (KO) cells differentiated on 2D and 3D culture formats were assessed by gene expression (normalized to *Actb* (encodes β-actin)). qPCR was run for reported early (**Panel A**; *Mmp14* and *Pdpn*) and intermediate (**Panel B**; *Phex* and *Dmp1*) markers of osteoblast (Obs) – osteocyte (Ocy) transition and subsequent differentiation. Columns not sharing a letter are significantly different via Tukey’s post-hoc test following applicable 3-Way (main effects: substrate, genotype, timepoint), p<0.05. Data is presented mean ± SD with data points for each column representing total n.

**Figure 5.**
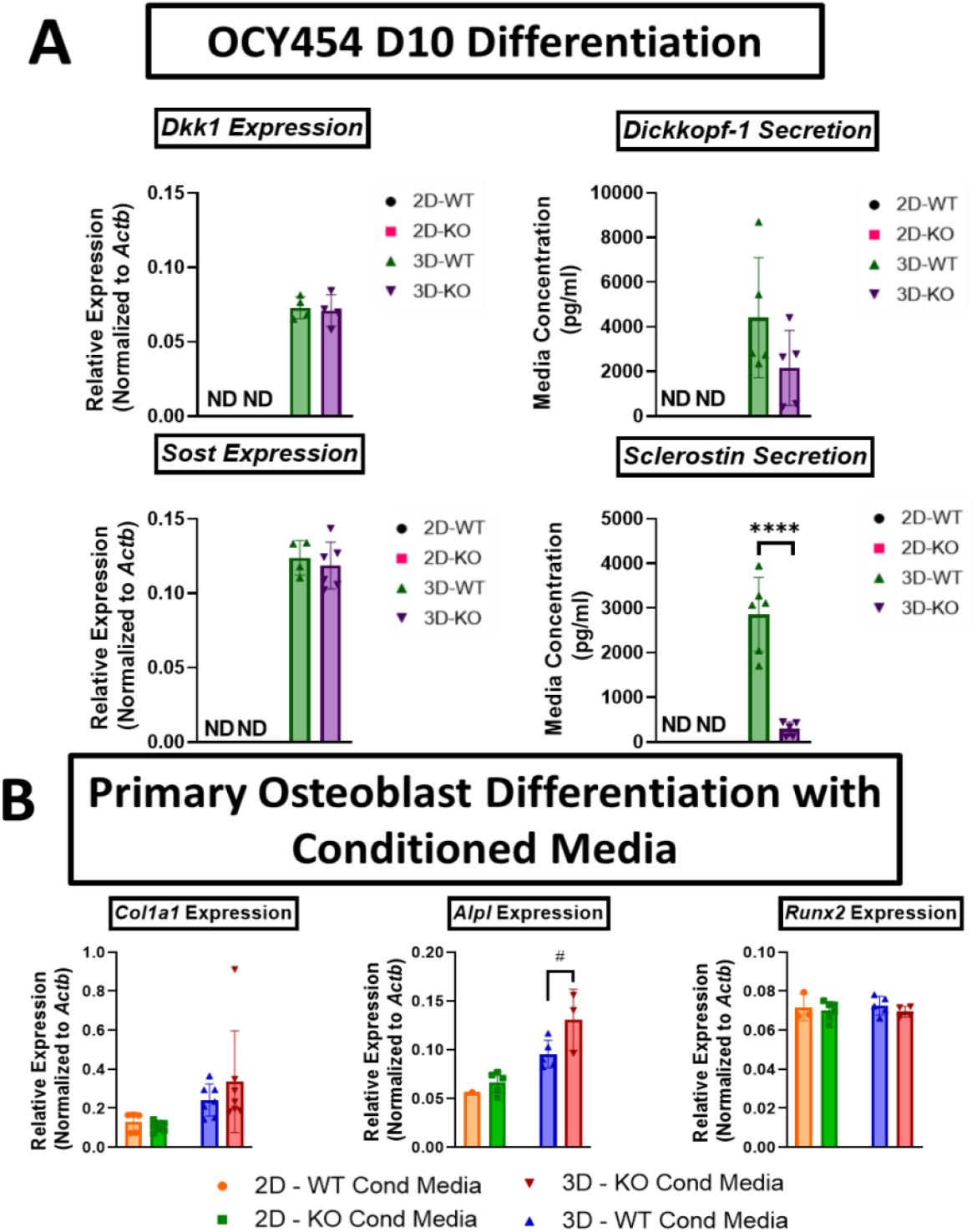
OCY454 Cx43 deficiency and cell culture format differentially regulated sclerostin secretion and osteoblastogenesis. Panel A. OCY454 osteocyte-like wildtype (WT) and Cx43-deficient (KO) differentiated for 10 days (D10) in 2D and 3D were assessed by gene expression (normalized to *Actb* (encodes β-actin)) for *Sost* and *Dkk1* expression and conditioned media by ELISA for sclerostin and dickkopf-1 secretion, respectively. ND = Not Detected (Outside Range of Assay). **** p < 0.00005; by unpaired t-test for 3D cell type (WT vs. KO). **Panel B:** Gene expression for osteoblast markers of differentiation (*Runx2, Alpl, Col1a1*) following 21 days of primary murine bone marrow stromal cell differentiation using conditioned media from D10 differentiation OCY454 cells. Primary bone marrow stromal cells from skeletally mature C57BL/6J mice showed trends (# p < 0.10) toward increased osteoblastic differentiation when treated with D10 conditioned media from 3D-KO versus 3D-WT OCY454 cells. **** p < 0.00005; # p < 0.10 by unpaired t-test for conditioned media type (WT vs. KO) within each cell culture format. Data is presented mean ± SD with data points for each column representing total n.

### Culture Substrate and Cx43 Deficiency Differentially Regulate Sclerostin Secretion and Osteoblastogenesis

Next, we examined how Cx43 deficiency and cell culture substrate differentially regulated OCY454 signaling to osteoblasts. To do this, we probed conditioned media of D10 differentiated OCY454 cells for DKK1 and sclerostin, key WNT/B-catenin inhibitors by which osteocytes regulate osteoblastogenesis.(6, 29) Measurement of sclerostin and DKK1 secretion into conditioned media by ELISA at D10 revealed significant differences in levels of protein secretion based on Cx43 deficiency and cell culture substrate. Similar to gene expression for *Dkk1* and *Sost*, all 2D cultured OCY454 cells (WT and KO) showed no detectable DKK1 and sclerostin protein in culture media (**Figure 5A**). Interestingly, while showing no significant difference between groups for *Sost* expression, 3D-KO cells displayed decreased levels of secreted sclerostin, relative to 3D-WT (**Figure 5A**; p < 0.0001). A similar trend was seen for DKK1 secretion (**Figure 5A;** p < 0.15), despite similar levels of *Dkk1* expression on samples at D10 between OCY454 WT and KO (**Figure 5A;** p > 0.05). To assess whether these changes in D10 conditioned media lead to altered osteoblast differentiation, primary bone marrow stromal cells were cultured in 50% conditioned media:50% standard OCY454 media for 21 days, and gene expression was assessed to determine osteoblastogenesis. Gene expression of osteoblast differentiation markers *Runx2, Alpl*, and *Col1a1* expression was not affected by conditioned media based on OCY454 Cx43 deficiency when the primary osteoblasts were cultured with 2D media (**Figure 5B**). However, *Alpl* expression trended (p < 0.10) upward when cultured with 3D-KO conditioned media compared to 3D-WT conditioned media, suggesting modest increased osteoblastogenesis signaling from 3D-KO OCY454 cells (**Figure 5B**). While 2D and 3D conditioned media were not directly compared to each other statistically at D10 due to large differences in cell count and thus potentially media protein levels, gene expression of mature osteoblast markers of mineralization and collagen secretion *Alpl* and *Col1a1* increased nearly 2-fold in 3D conditioned media samples compared to 2D counterparts. These results suggest secreted factors, other than DKK1 and sclerostin, from more mature OCY454 osteocytes cultured on 3D versus 2D, may further promote osteoblast differentiation.

### Culture Substrate and Cx43 Deficiency Differentially Regulated RANKL/OPG Ratio and Osteoclastogenesis

Next, we investigated how Cx43 deficiency and cell culture substrate may differently regulate OCY454 signaling to osteoclasts. To do this, we assessed gene expression of key osteoclastogenic molecules produced by osteocytes, *Tnfsf11* and *Tnfrsf11b* (encodes RANKL and OPG, respectively) from OCY454 cells at D10 of differentiation.(30) Significant main effects of both genotype (p < 0.0001), substrate (p = 0.0252) and interaction (p < 0.0001) were found at D10 for the ratio of *Tnfsf11/Tnfrsf11b* gene expression by OCY454 cells. For example, the ratio of *Tnfsf11/Tnfrsf11b* gene expression was significantly increased in OCY454 cells cultured on 2D (+7.94%) and in KO cells versus WT cells (+23.2%; **Figure 6A**) overall. Due to a significant interaction effect, 2D-KO cells displayed significantly higher *Tnfsf11/Tnfrsf11b* expression compared to 2D-WT, but this was not found to be significantly different when directly comparing 3D–WT and 3D-KO samples (**Figure 6A**). In addition, KO cells displayed higher *Tnfsf11/Tnfrsf11b* expression on 2D compared to 3D, whereas the converse was true for WT cells. Overall, this suggests a complex relationship between cell culture substrate and Cx43 deficiency in regard to osteoclastogenic signaling, with increased *Tnfsf11/Tnfrsf11b* expression in KO cells, but only those grown on 2D compared to 3D. TRAP staining and gene expression for the bone resorption enzyme *Ctsk*, were used to determine relative osteoclastogenesis of primary bone marrow stromal cells (BMSCs) when cultured for 6 days with OCY454 conditioned media supplemented with RANKL (2.5 ng/ml) and M-CSF (25ng/ml). Contrary to *Tnfsf11/Tnfrsf11b* expression in OCY454 cells, osteoclast size was significantly increased in cells treated with 3D-KO conditioned media compared to that of 3D-WT, and this trend was also observed when using 2D conditioned media (**Figure 6B**; p = 0.16). *Ctsk* expression was also significantly increased in primary BMSCs with 2D-KO conditioned media compared to that of 2D-WT (**Figure 6B**; p <0.005). While not statistically compared, BMSCs treated with 3D conditioned media induced greater osteoclast formation as evidenced by increased osteoclast size and *Ctsk* expression than those treated with 2D conditioned media for both WT and KO genotypes.

**Figure 6.**
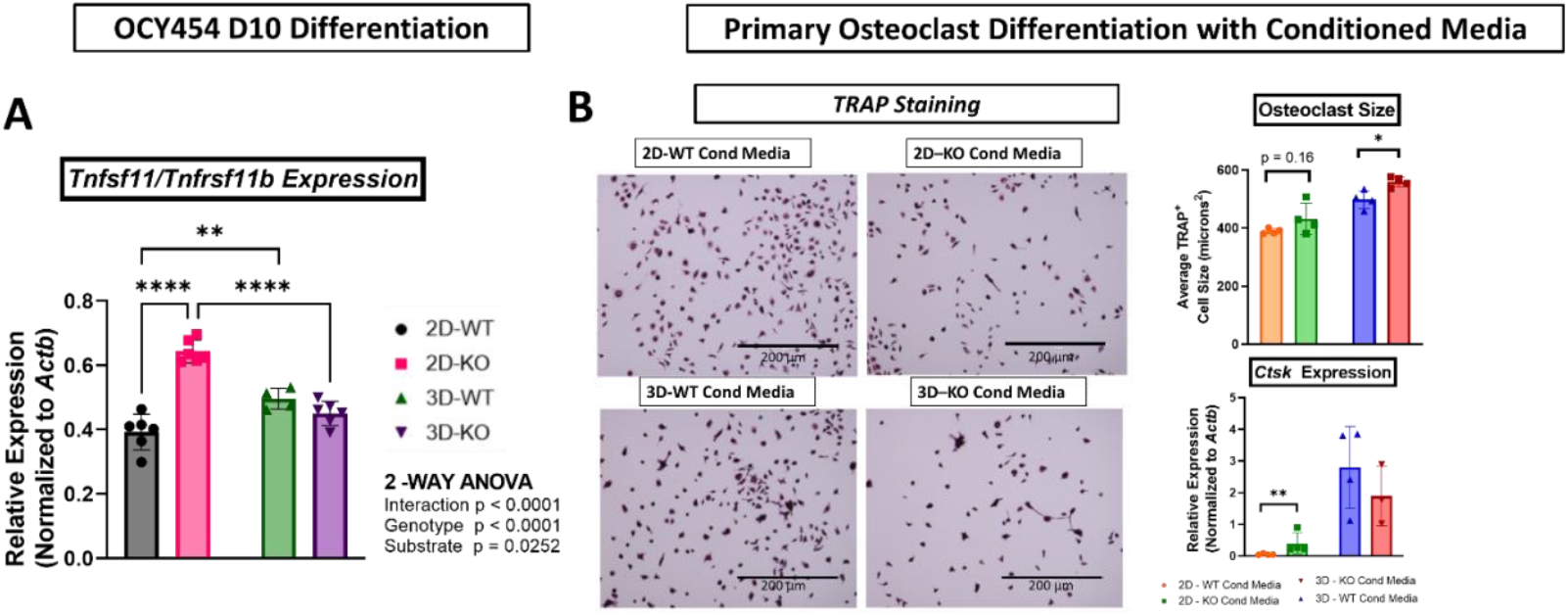
OCY454 Cx43 deficiency and cell culture format differentially regulate TNFSF11/TNFRSF11b expression and osteoclastogenesis. Panel A. OCY454 osteocyte-like wildtype (WT) and Cx43-deficient (KO) cells cultured on 2D and 3D formats were assessed at day 10 of differentiation (D10) by gene expression (normalized to *Actb* (encodes β-actin)) for *Tnfsf11/Tnfrsf11b* expression (encodes RANKL/OPG), respectively. **Panel B:** TRAP staining and gene expression for osteoclast resorption enzyme *CTSK* (encodes cathepsin K) following 6 days of primary murine bone marrow stromal cell differentiation using conditioned media from D10 differentiation OCY454 cells supplemented with RANKL (5ng/ml) and M-CSF (25ng/ml). Overall, conditioned media KO cells, especially those grown in 3D, appeared to synergistically promote osteoclastic differentiation of bone marrow stromal cells from skeletally mature C57BL/6J mice. **** p < 0.00005; ** p <0.005; * p < 0.05 by Tukey’s post-hoc test following unpaired t-test for conditioned media type (WT vs. KO) within each cell culture format (2D vs. 3D). Data is presented mean ± SD with data points for each column

## Discussion

In this study, we studied both the effects of Cx43 deficiency and cell culture substrate on cell autonomous differentiation and signaling to bone effector cells (osteoblasts and osteoclasts) in the immortalized osteocytic OCY454 cell line. Our results show combined effects of both Cx43 deficiency and cell culture substrate on OCY454 differentiation and signaling to promote osteoblastogenesis and osteoclastogenesis. Conditioned media from KO cells increased both primary cell osteoblast and osteoclast differentiation, compared to conditioned media from WT cells. This same effect was observed when comparing media taken from cells cultured on 3D versus 2D, with media from 3D cultured OCY454 cells promoting increased osteoblast and osteoclast differentiation of primary BMSCs. These data support our first hypothesis that Cx43 deficiency in osteocytes enhances paracrine signaling to promote increased osteoblast and osteoclast differentiation. These results indicate the increased bone remodeling seen in Cx43 deficient bone *in vivo*, may be due to cell autonomous increases in osteocyte paracrine signaling.(13, 14) However, since OCY454 sclerostin and *Tnfsf11/Tnfrsf11b* expression changes didn’t always follow trends in osteoblastogenesis and osteoclastogenesis, it’s unlikely the Cx43-deficient phenotype is solely working through these molecules. In addition, we observed decreased proliferation and increased differentiation of OCY454 cells on 3D compared to 2D. However, regardless of cell culture substrate, Cx43 deficiency did not appear to dramatically delay or inhibit OCY454 cell differentiation, thereby refuting our second hypothesis. Overall, these results suggest Cx43 modulates OCY454 paracrine signaling, while decreased cell count and increased expression of osteocyte maturation genes on 3D Alvetex cultured samples suggest OCY454 cell differentiation is largely influenced by culture substrate over Cx43 deficiency in our study.

Analysis of OCY454 cell count by dsDNA content following seeding demonstrated differing cell dynamics due to Cx43 deficiency and cell culture substrate. We observed higher rates of cell proliferation on 2D versus 3D (evidenced by increased cell count and *Ccnd1* expression), (**Figure 3A**). Since both 3D Alvetex and 2D tissue culture plastic were made of the same material (0.8 mg/ml Collagen 1 coated polystyrene), it’s more likely physical interactions and potentially reduced growth area in the Alvetex, contributed partially to this phenotype. In contrast, on Alvetex, cell counts reached their maximum by day 6 of differentiation, with significantly more cells in 3D-WT versus 3D-KO (50% more) despite equal seeding densities (**Figure 3A**). This greater number of 3D–WT versus 3D–KO may be due to both differential rates of cell proliferation and apoptosis that weren’t fully captured by our analysis. Previous reports have demonstrated normal to slightly increased rates of cell proliferation and elevated apoptosis due to loss of Cx43 in osteoblasts and osteocytes *in vivo* and *in vitro*.(13, 22, 31–33) In contrast to these data, only *Ccnd1* (proliferation) and not *Ddit3 (apoptosis)* expression was elevated in 3D– WT versus 3D–KO cells (trending increase p < 0.10 at D3; **Figure 3B**). These conflicting results may be due to the different immortalized cell type used in our study (OCY454 versus MLO-Y4), starting seeding densities or culturing cells on the 3D substrates. Thus, our results suggest Alvetex decreases OCY454 proliferation, which may be further decreased with Cx43 deficiency.

Cell culture substrate was more impactful on OCY454 differentiation than Cx43 deficiency. We saw significantly increased expression of early (*Dmp1, Phex*) and late (*Sost, Dkk1*) differentiation markers on 3D versus 2D as early as D6 (**Figure 4B**) and D10 (**Figure 5A**), respectively. However, at these same timepoints, there were no significant pair-wise changes in osteocyte gene expression due to Cx43 deficiency on either cell culture substrate. These results suggest in OCY454 cells, cell culture substrate affects osteocyte terminal differentiation while Cx43 deficiency does not. The enhanced OCY454 osteocyte maturation on 3D Alvetex versus 2D tissue culture plastic has been previously reported.(19) Furthermore, a recent study demonstrated a 10-40 fold increase in *Sost* expression in MLO-Y4 cells cultured in 3D versus 2D Collagen1-Hydroxyapatite gels.(34) Thus, similar to our study, substrate dimensionality clearly influenced osteocytic differentiation, however the underlying mechanism remains unknown.

The enhanced differentiation may be partially due to Alvetex induced increases in WNT/β-catenin pathway activation. For example, osteocytic WNT activation in mice by a nondegradable β-catenin transgene increases osteocytic markers of differentiation, viability and bone turnover.(35) In our study, WT and KO cells on 2D TCPS by D10 showed decreased expression of mature osteocyte markers (*Dmp1, Dkk1, Sost;* **Figure 4B-5A**) and Wnt/β-catenin canonical pathway target genes (*Ctnnb1, Tnfrsf11b;* **Supplemental Figure S3**) compared to 3D Alvetex, suggesting increased differentiation concomitant with increased WNT/β-catenin canonical pathway activation on 3D Alvetex. However, unlike previous results demonstrating Cx43 deficiency decreases osteocytic gene expression of *Dmp1* and *Sost* expression by over 80% in ID3-SWG cells cultured in 2D, we saw little differences in OCY454 *Dmp1* and *Sost* gene expression due to Cx43 deficiency.(8) The discrepancy may be due to a block in osteoblast-osteocyte transition as noted by Hua et al, as ID3-SWG cells first start off as osteoblasts compared to early osteocytes in the OCY454 cell line.(8) In addition, it may be because the Alvetex was such a potent inducer of differentiation of both WT and KO cells, that Cx43 effects on differentiation were diminished. Furthermore, the lack of strong Cx43 deficient interactions on differentiation could be due to the relative lower concentration of Cx43 protein reported to be produced by OCY454 cells compared to ID3-SWG cells and MLO-Y4 cells as reported by others and seen by us (**Supplemental Figure S1**).(8, 19) Cx43 deficiency did not impact osteocyte differentiation but did impair GJIC (**Figure 2B**), suggesting GJIC may not be a strong prerequisite for osteocyte differentiation, as it has been previously reported in osteoblastic cell lines.(36–38) These mechanisms of enhanced osteocyte differentiation on Alvetex and other 3D culture models warrant further study.

Despite minor effects of Cx43 deficiency on osteocyte differentiation, we saw more pronounced effects of Cx43 deletion on osteocyte conditioned media based signaling to effector cell populations. Conditioned media experiments demonstrated Cx43 deficiency and cell culture substrate regulated secretion of the WNT/B-catenin inhibitor sclerostin (and to a lesser extent DKK1) and resultant osteoblastogenesis. For example, although there were no significant changes in *Sost* or *Dkk1* gene expression between 3D-KO and 3D-WT samples at D10, we measured a ∼90% reduction in secreted sclerostin and ∼50% reduction in secreted DKK1 due to Cx43 deficiency. Importantly, the decreased sclerostin secretion (**Figure 5A**) but not DKK1 was much greater than the 50% reduction in cell population seen at D10 between 3D–KO and 3D–WT cells (**Figure 3A**). In all, these results suggest on a per cell basis, Cx43 deficiency may be affecting post-transcriptional modification and/or secretion of sclerostin from osteocytes.

Although outside the scope of this manuscript, reported mechanisms regulating sclerostin protein production include lysosomal degradation, focal adhesion/integrin signaling, and oxygen sensing and should be investigated further with our OCY454 Cx43 deficient cells.(39, 40) Examining primary BMSC cell culture, we saw trends toward increased late-stage osteoblast differentiation (*Col1a1, Alp* expression) when using 3D–KO versus 3D–WT conditioned media, although this was not significant. Potentially enhanced osteoblast differentiation from KO OCY454 cells is in line with *in vivo* Cx43 deficient cortical bone showing localized increased periosteal formation by dynamic labeling.(13) Although primary osteoblasts from 2D and 3D conditioned media samples weren’t directly compared statistically (due to vastly different cell and protein counts), 3D conditioned media on a volume/volume basis enhanced late-stage osteoblast differentiation (via *Col1a1, Alp* expression) compared to 2D samples overall. This is surprising, considering 3D Alvetex dramatically increased secreted sclerostin, a known inhibitor of osteoblast formation, in both WT and Cx43 KO cells compared to 2D TCPS. Therefore, 3D Alvetex may alter the milieu of other secreted factors (not tested here) that better support osteoblast differentiation overall compared to more immature pre-osteocytes grown on 2D TCPS.(41, 42)

Conditioned media osteoclast experiments also demonstrated an interaction between Cx43 deficiency and cell culture substrate in regulating *Tnfsf11/Tnfrsf11b* (encodes RANKL/OPG) expression and resultant osteoclastogenesis. For example, we saw significantly higher expression of *Tnfsf11/Tnfrsf11b* in 2D–KO versus 2D–WT OCY454 cells (**Figure 6A**) with higher rates of osteoclast differentiation in primary BMSCs cultured with 2D-KO conditioned media versus 2D–WT conditioned. This increased osteoclast differentiation was marked by increased trends in osteoclast size and significantly elevated *Ctsk* expression (**Figure 6B**). In contrast, there were no significant differences in *Tnfsf11/Tnfrsf11b* in 3D–KO versus 3D–WT OCY454 cells (**Figure 6A**), despite significant differences in osteoclast size when primary BMSCs are cultured with conditioned media from 3D–KO versus 3D–WT OCY454 cells (**Figure 6B**). Qualitatively, osteoclast size and *Ctsk* expression were greater in BMSCs cultured with OCY454 conditioned media from 3D versus 2D, despite *Tnfsf11/Tnfrsf11b* expression being greater overall in 2D cultures. These results suggest OCY454 KO cells secrete more factors promoting osteoclastogenesis compared to OCY454 WT cells, which appear to correlate with *Tnfsf11/Tnfrsf11b* expression in less mature 2D grown OCY454 osteocytes, but not more mature 3D grown OCY454 osteocytes. A limitation of our study is that we evaluated *Tnfsf11/Tnfrsf11b* expression and not RANKL/OPG secretion as we were unable to detect significant amounts of RANKL in OCY454 conditioned media. Therefore *Tnfsf11/Tnfrsf11b* expression may not accurately represent RANKL/OPG signaling from OCY454 cells to primary BMSCs. This is supported by the fact 3D–WT versus 3D–KO demonstrated no significant changes in *Tnfsf11/Tnfrsf11b* expression (**Figure 6A**) despite profound changes in BMSC osteoclastogenesis (**Figure 6B**). It’s also possible other known osteoclastogenic cytokines and molecules secreted by osteocytes, but not directly assayed here, such as extracellular ATP, IL-6 and TNF-α(43–45), are also influencing our osteoclast assays Nonetheless, the increased osteoclastogenesis seen in our study when utilizing conditioned media from OCY454 KO cells is consistent with previous results in Cx43 deficient MLO-Y4 cells and ID3-SWG cells and *in vivo* Cx43 deficient bone.(8, 22, 43) Therefore, our results further give strong support for the cell autonomous role Cx43 plays in regulating osteocyte control of osteoclastogenesis.(14)

In summary, our results demonstrate Cx43 deficiency and the cell culture environment combinatorically influence osteocyte biology, which previously has not been explored simultaneously. Our study suggests osteocytic Cx43 deficiency, when using the more mature osteocyte-like OCY454 cell line, demonstrates Cx43 is dispensable for osteocyte differentiation but greatly impacts osteocytic signaling to support osteoblastogenesis and osteoclastogenesis. Furthermore, we demonstrate the cell culture substrate, as expected, greatly impacts the findings of osteocyte biology experiments. Utilization of 3D Alvetex over 2D TCPS led to less cellular proliferation, greater differentiation and generated a more mature osteocytic phenotype, marked by sclerostin and DKK1 secretion, which were altered due to Cx43 deficiency. This exciting result should be further explored to determine how substrate sensing and Cx43 may regulate sclerostin secretion. Furthermore, conditioned media from OCY454 cells cultured on 3D Alvetex were better able to differentiate primary BMSCs into osteoblasts and osteoclasts than OCY454 cells grown on 2D TCPS. However, this was surprising considering conditioned media from 3D Alvetex had higher sclerostin secretion and lower *Tnfsf11/Tnfrsf11b* expression. This suggests other osteocyte secreted factors in the 3D Alvetex versus 2D TCPS conditioned media may also be at work and should be explored in future directions.

Nonetheless, Cx43 deficiency in both 2D TCPS and 3D Alvetex cultures caused increased osteoblastogenesis and osteoclastogenesis, suggesting increased bone turnover and remodeling due to decreased GJIC, which is supported by *in vivo* studies.(13, 14) Therefore, future work studying osteocyte biology and signaling to other cell types, should utilize osteocytes grown in a 3D microenvironment to better mimic *in vivo* experiments and explore mechanisms. In all, modulation of osteocytic Cx43 is a promising target to cell autonomously control bone mass and strength via osteocytic coordination of bone arms of bone remodeling.

## Acknowledgements

We would like to thank Joshua Cohen, M.D. and Kayla Scott of the Boyan Lab of Musculoskeletal Research and Innovation (LMRI) at VCU BME for histological and imaging assistance. We would like to thank Shirley M Taylor, Ph.D. at Biological Macromolecule Shared Resource (supported, in part, with funding from NIH-NCI Cancer Center Support Grant P30 CA016059), who helped to create OCY454 Cx43 deficient cell line. This work was supported by the VCU College of Engineering Foundation Research Endowment, NASA (Grant #: 80NSSC18KK1473) and the NIH (Grant # R01AR068132). Drs. Evan Buettmann and Michael Friedman were supported by the Translational Research Institute for Space Health (TRISH) through Cooperative Agreement with NASA (NNX16AO69A).

## Author Roles

Authors’ Roles: Study design: HJD, YZ, MAF, GAH, EGB, GMG. Study conduct: GAH, EGB, JAD, SNC, YZ, GMG Data collection: GAH, EGB, JAD, SNC, GMG, YZ. Data analysis: GAH, EGB, JAD, SNC, GMG, YZ. Data interpretation: HJD, YZ, MAF, GAH, EGB, JAD, GMG. Drafting manuscript: GAH, EGB, JAD. Revising manuscript: GAH, EGB, GAG, HJD, MAF, YZ. Approving final version of manuscript: All authors take responsibility for integrity of data analysis.

