## Supplemental Figure 1 for "Connexin 43 and Cell Culture Substrate Differentially Regulate OCY454 Osteocytic Differentiation and Signaling to Primary Bone Cells"

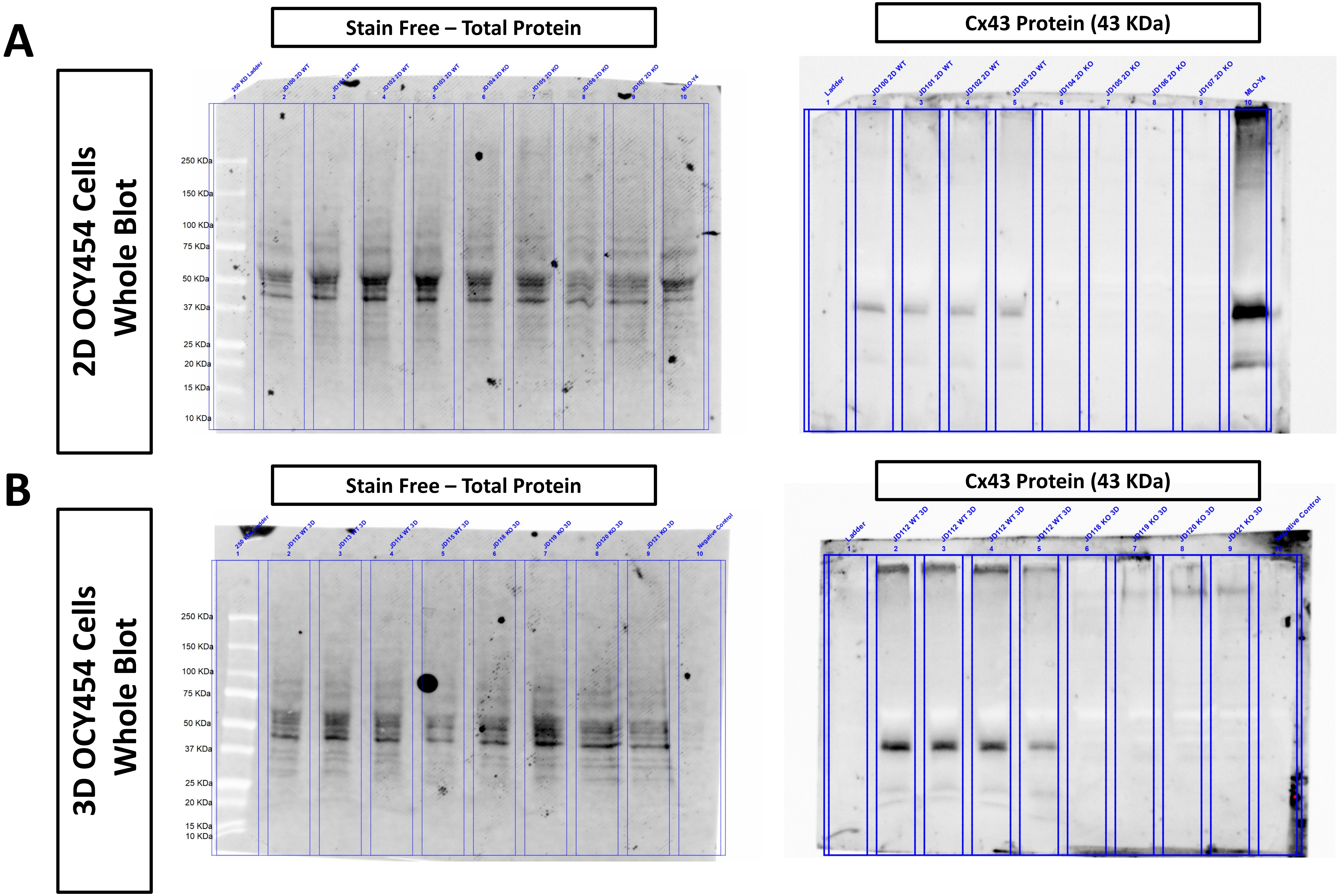


**Supplemental Figure S1. Validation of Cx43 Deficiency in OCY454 Cells. A:**Total cell lysate from OCY454 osteocyte-like wildtype (OCY454 WT) and MLO-Y4 cells differentiated for 10 days on TCPS were loaded at equal protein concentrations (50 ug) onto gels showed robust Cx43 protein staining (Sigma C6219 1: 8000) on western blot compared to OCY454 Cx43 deficient cells (KO). **B:** Similarly, total cell lysate from osteocyte-like wildtype (WT) cells cultured on Alvetex for 10 days and loaded at equal protein concentrations (50 ug) onto gels showed robust Cx43 protein staining on western blot, with minimal Cx43 protein detected in Alvetex cultured OCY454 Cx43 deficient (KO).
