## Supplemental Figure 2 for "Connexin 43 and Cell Culture Substrate Differentially Regulate OCY454 Osteocytic Differentiation and Signaling to Primary Bone Cells"

### WNT/β-catenin Target Genes

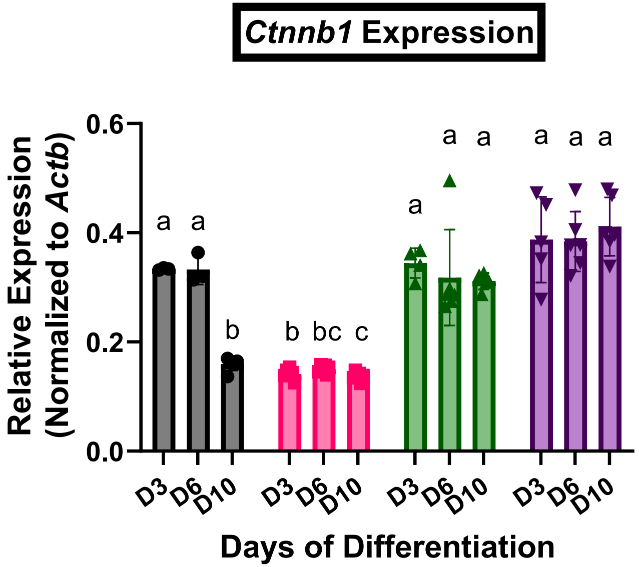


**3-WAY ANOVA Table**
Timepoint $p < 0.0001$
Genotype $p < 0.0001$
Substrate $p < 0.0001$
Timepoint X Genotype $p < 0.0001$
Timepoint X Substrate $p < 0.0001$
Genotype X Substrate $p < 0.0001$
Timepoint X Genotype X Substrate $p < 0.0001$

● 2D-WT
■ 2D-KO
▲ 3D-WT
▼ 3D-KO

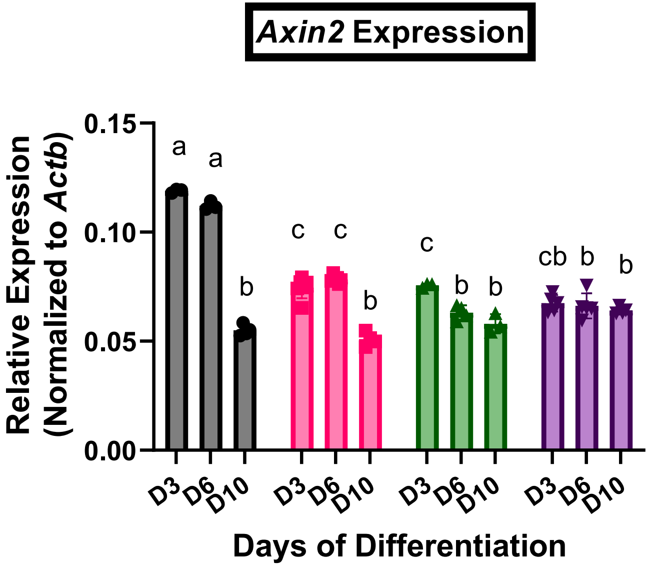


**3-WAY ANOVA Table**
Timepoint $p < 0.0001$
Genotype $p < 0.0001$
Substrate $p < 0.0001$
Timepoint X Genotype $p < 0.0001$
Timepoint X Substrate $p < 0.0001$
Genotype X Substrate $p < 0.0001$
Timepoint X Genotype X Substrate $p = 0.4752$

● 2D-WT
■ 2D-KO
▲ 3D-WT
▼ 3D-KO

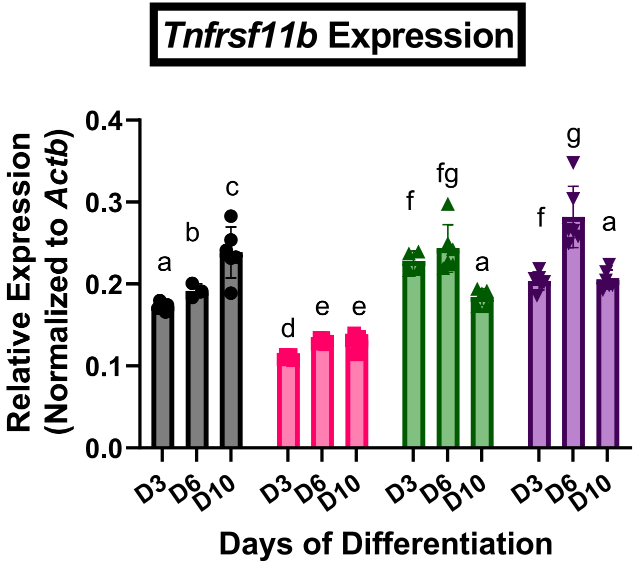


**3-WAY ANOVA Table**
Timepoint $p < 0.0001$
Genotype $p < 0.0001$
Substrate $p < 0.0001$
Timepoint X Genotype $p = 0.0008$
Timepoint X Substrate $p < 0.0001$
Genotype X Substrate $p < 0.0001$
Timepoint X Genotype X Substrate $p = 0.0206$

● 2D-WT
■ 2D-KO
▲ 3D-WT
▼ 3D-KO

**Supplemental Figure S2. OCY454 Cx43 deficiency and culture format differentially regulate WNT/β-catenin target gene expression.** Changes in WNT/β-catenin target OCY454 osteocyte-like wildtype (WT) and Cx43-deficient (KO) cells on 2D (TCPS) and 3D (Alvetex) cell culture formats were assessed by gene expression (normalized to *Actb* (encodes β-actin)). qPCR was run for reported target genes *Bcat* (encode β-catenin), *Axin2* (encodes Axis Inhibition Protein 2), *Tnfrsf11b* (encodes Osteoprotegerin), known to have increased transcription with WNT/β-catenin canonical pathway activation. By D3, 2D-WT had elevated expression of *Bcat*, *Axin2* and *Tnfrsf11b* compared to 2D-KO cells on TCPS only, suggesting a delay in WNT/β -catenin activation with Cx43 deficiency in 2D. By later (D10) timepoints of differentiation, 2D-WT still had significantly increased markers of WNT/β-catenin canonical pathway activation compared to 2D-KO Cells (*Bcat and Tnfrsf11b).* In contrast, 3D-WT and 3D-KO showed similar minor attenuation in WNT/β-catenin canonical pathway target genes over time compared to 2D (regardless of genotype), suggesting differential WNT/β -catenin pathway activation on Alvetex. Columns not sharing a letter are significantly different via Tukey's post-hoc test following applicable 3-Way (main effects: substrate, genotype, timepoint). Data is presented mean ± SD with data points for each column representing total n.
