## Supplemental Table 1 for "Connexin 43 and Cell Culture Substrate Differentially Regulate OCY454 Osteocytic Differentiation and Signaling to Primary Bone Cells"

**Supplemental Table 1. Standard Media Types for OCY454 Cell Differentiation Assays and Primary Osteoblast and Osteoclast Conditioned Media Experiments**

| **Media Name** | **Abbreviation** | **Base Ingredients** | **Added Supplements** |
| --- | --- | --- | --- |
| Standard OCY454 Media | Standard OCY454 αMEM | 10% FBS; 2% Penn/Strep | 2.2 g/L Sodium Bicarbonate |
| Osteoblast Positive Control Media | OB Diff Media | 10% FBS; 2% Penn/Strep | 2.2 g/L Sodium Bicarbonate; 50 μg/mL L-ascorbic acid-2-phosphate; 10 mM β-Glycerophosphate |
| Osteoclast Differentiation Media | OC Diff Media | 10% FBS; 2% Penn/Strep | 2.2 g/L Sodium Bicarbonate; 25 ng/ml M-CSF; 5 ng/mL RANKL |
| Osteoclast Negative Control Media | OC NC Media | 10% FBS; 2% Penn/Strep | 2.2 g/L Sodium Bicarbonate; 25 ng//ml M-CSF |
| Osteoclast Positive Control Media | OC PC Media | 10% FBS; 2% Penn/Strep | 2.2 g/L Sodium Bicarbonate; 25 ng/mL M-CSF; 25 ng/mL RANKL |
