## Supplemental Table 2 for "Connexin 43 and Cell Culture Substrate Differentially Regulate OCY454 Osteocytic Differentiation and Signaling to Primary Bone Cells"

**Supplemental Table 2. Markers of Cell Proliferation, Osteocyte Differentiation, and Bone Remodeling**

| **Gene Symbol** | **Gene Name** | **Gene Function** | **Assay ID:** | **Ensemble ID (Splice Variants Targeted):** | **Chromosome Location:** |
| --- | --- | --- | --- | --- | --- |
| *Ccnd1* | Cyclin D1 | Activity is required for cell cycle G1/S transition | qMmuCID0023518 | [ENSMUSG00000070348](http://www.ensembl.org/id/ENSMUSG00000070348) | 7:144932712-144934061 |
| *Ddit3* | DNA Damage Inducible Transcript 3 | Activated by endoplasmic reticulum stress and promotes apoptosis | qMmuCID0005629 | [ENSMUSG00000025408](http://www.ensembl.org/id/ENSMUSG00000025408) | 10:127290791-127293751 |
| *Mmp14* | Matrix Metallopeptidase 14 | Expressed early during osteoblast-osteocyte transition | qMmuCID0006120 | [ENSMUSG00000000957](http://www.ensembl.org/id/ENSMUSG00000000957) | 14:54437681-54438722 |
| *Pdpn* | Podoplanin | Expressed early during osteoblast-osteocyte transition | qMmuCED0048126 | [ENSMUSG00000028583](http://www.ensembl.org/id/ENSMUSG00000028583) | 4:143273912-143274041 |
| *Dmp1* | Dentin Matrix Acidic Phosphoprotein 1 | Regulates osteocyte maturation and phosphate metabolism | qMmuCID0008852 | [ENSMUSG00000029307](http://www.ensembl.org/id/ENSMUSG00000029307) | 5:104210102-104211822 |
| *Dkk1* | Dickkopf-Related Protein 1 | Produced in mature osteoblasts/osteocytes, negative regulator of WNT pathway | qMmuCID0020261 | ENSMUST00000025803 | 19:30547563-30548803 |
| *Sost* | Sclerostin | Produced in mature osteocytes, negative regulator of WNT pathway | qMmuCID0020701 | [ENSMUSG00000001494](http://www.ensembl.org/id/ENSMUSG00000001494) | 11:101964208-101966847 |
| *Tnfsf11* | TNF Superfamily Member 11 (RANKL) | Key factor for osteoclast differentiation and activation | qMmuCID0026078 | [ENSMUSG00000022015](http://www.ensembl.org/id/ENSMUSG00000022015) | 14:78284239-78299916 |
| *Tnfrsf11b* | TNF Receptor Superfamily Member 11b (Osteoprotegerin) | Negative regulator of bone resorption; RANKL inhibitor | qMmuCID0027158 | [ENSMUSG00000063727](http://www.ensembl.org/id/ENSMUSG00000063727) | 15:54254138-54256042 |
