## Supplemental Table 3 for "Connexin 43 and Cell Culture Substrate Differentially Regulate OCY454 Osteocytic Differentiation and Signaling to Primary Bone Cells"

**Supplemental Table 3. Markers of Osteoblast and Osteoclast Differentiation**

| **Gene Symbol** | **Gene Name** | **Gene Function** | **Assay ID:** | **Ensemble ID (Splice Variants Targeted):** | **Chromosome Location:** |
| --- | --- | --- | --- | --- | --- |
| *Ctsk* | Cathepsin K | Cysteine proteinase involved in bone resorption | qMmuCID0022824 | [ENSMUSG00000028111](http://www.ensembl.org/id/ENSMUSG00000028111) | 3:95506692-95508910 |
| *Alpl* | Alkaline Phosphatase | Enzyme regulating bone mineralization | qMmuCID0006482 | [ENSMUSG00000028766](http://www.ensembl.org/id/ENSMUSG00000028766) | 4:137755656-137758687 |
| *Runx2* | RUNX2 Family Transcription Factor 2 | Transcription factor necessary for osteoblast differentiation and gene expression | qMmuCED0049270 | [ENSMUSG00000039153](http://www.ensembl.org/id/ENSMUSG00000039153) | 17:44724789-44735235 |
| *Col1a1* | Collagen Type 1, Alpha 1 | Procollagen Alpha Chain of Collagen Trimer Molecule | qMmuCED0044222 | [ENSMUSG00000001506](http://www.ensembl.org/id/ENSMUSG00000001506) | 11:94950400-94950515 |
